# Nucleosome-scale p53–hormone receptor grammar distributed by Alu elements

**DOI:** 10.64898/2026.09.26.754600

**Authors:** Xi Fu, Qingyuan Cai, Pranay Satya, Raul Rabadan, Arnold J. Levine

**Author notes:** Equal contribution (Xi Fu, Qingyuan Cai).

## Abstract

The tumor suppressor p53 and the transcription factors estrogen receptor and androgen receptor regulate growth in a sex-dependent hormone-responsive manner, as in breast and prostate tumors and muscle tissue. Using a model of transcriptional regulation, we investigated co-regulation between p53 and hormone receptors in different cellular contexts and highlight an enrichment in mammary and muscle cell types. Sequence, structural and epigenomic evidence demonstrate a broadly distributed functional 73-bp p53-estrogen receptor grammar spanning nucleosome entry/exit to dyad, while p53-androgen receptor grammar spans 146 bp. At a functional breast cancer enhancer regulating Cyclin D, varied spacing led to lower predicted p53 binding. Genome-wide analysis found a major role of Alu sequences in distributing this grammar, in particular in mTORC1 pathway genes, with an evolutionary association of primate body-size sexual dimorphism and muscle aging.

## Introduction

p53 is a conserved stress-responsive regulator of genome integrity, cell-cycle arrest, and cell-fate decisions (*1*), whereas estrogen receptor (ER) and androgen receptor (AR) translate endocrine signals into cell-type transcription programs governing development, metabolism, and reproduction. ER drives enhancer activity and proliferative programs in breast tumors (*2*). Clinically, TP53 mutations are associated with primary endocrine resistance in ER-positive breast cancer and are enriched in subsets of metastatic tumors that progress despite combined endocrine and CDK4/6-targeted therapy (*3, 4*). Meanwhile, AR drives growth and nutrient-responsive programs in hormone-responsive tissues (*5, 6*), such as muscle and prostate tumors. Although p53 can cooperate with ER and AR in chromatin regulation (*5, 7, 8*), these factors belong to distinct families with different DNA-binding domains. How their binding sites are spatially arranged to support cooperation, and how such a grammar can recur across the genome, remain unclear.

Transposable elements have contributed to the expansion of p53 binding sites in primate genomes (*9*). Their repeated sequence architecture could distribute not only individual sites but also their spacing and orientation relative to hormone-receptor motifs. This raises the question of whether repeats carry p53–hormone receptor (HR) grammars that support co-regulation in distinct cellular contexts. We therefore investigated the cellular contexts, structural organization, regulatory implications, and evolutionary origins of these grammars or architectures in a cell.

## Results

### Nucleosome-scale grammar for p53 and hormone receptors

To identify cellular contexts in which p53-HR coregulation may occur, we applied General Expression Transformer (GET) (*10*), a foundation model pretrained on a single-cell atlas of chromatin accessibility and transcriptomes of human primary cell types (Fig. 1A). p53-ER coregulation is enriched in fetal skeletal myocytes (probably corresponding to estrogen-related receptors (*11*)), club cells (*12*), and mammary luminal epithelial cells (Fig. 1B, fig. S1B), whereas AR–p53 coregulation is enriched in fetal skeletal satellite cells and vascular- or smooth-muscle-related cells (Fig. 1C).

**Fig. 1.**
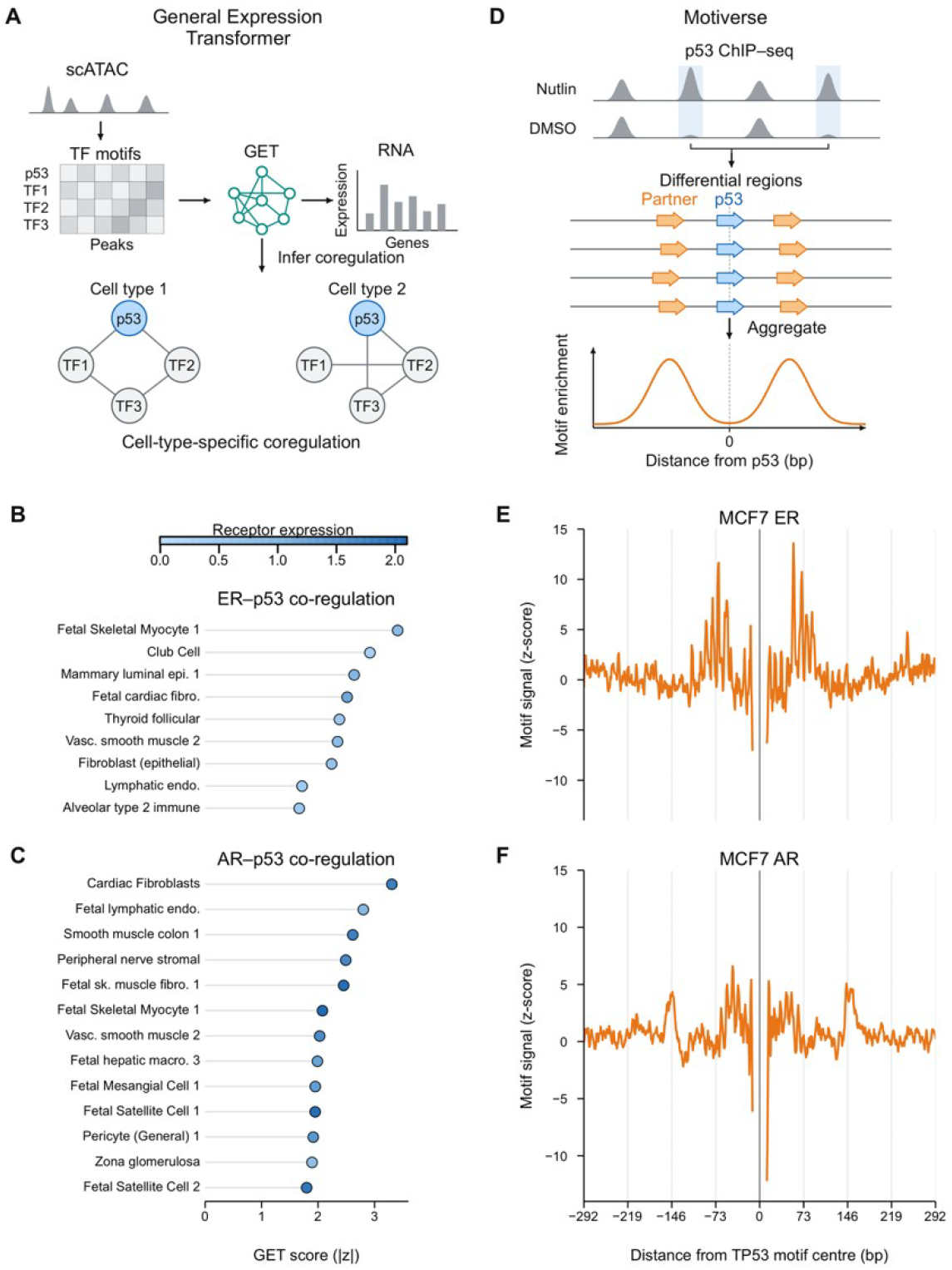
Cell context and spacing grammar of p53 and hormone receptors. **(A)** General Expression Transformer (GET) uses scATAC-derived TF motif-by-peak inputs to model cell-type-specific RNA expression and infer corresponding TF coregulation. **(B and C)** In silico study of p53-ER and p53-AR coregulation: GET coregulation scores for p53 with ER (B) or AR (C) across cell types; point color denotes the predicted expression of the corresponding receptor. **(D)** Motiverse algorithm aggregates position-specific partner motif profiles around p53 motifs within differential p53 ChIP-seq regions between nutlin and DMSO treatment. **(E and F)** Motiverse motif-position profiles for ER-associated (E) and AR-associated (F) motifs around 40,616 p53 motifs in nutlin-treated MCF7 TP53 ChIP-seq peaks. Motif signal is expressed as a z-score relative to background score on −500 to −293 bp and +293 to +500 bp from the p53 motif.

These transcription factors have been shown to cooperate as homotypic (*13, 14*) or heterotypic dimers (*15*), facilitated by DNA (*16, 17*), or guided by nucleosome fiber topology (*18*). To further dissect which category p53-HR might be in, we developed Motiverse, a GPU-accelerated algorithm that can perform one-to-all motif-pair spacing analyses across genome-wide regulatory regions in minutes (Fig. 1D, Supplementary Materials). Applying Motiverse to nutlin-induced p53 ChIP–seq peaks in MCF7 breast cancer cells, we identified an enrichment of ER motifs 73 bp away from the p53 binding site (Fig. 1E, fig. S1D). By contrast, the AR-associated profile showed an enrichment near ±146 bp, together with a small enrichment near 50 bp relative to the p53 anchor (Fig. 1F, fig. S1D). These numbers directed us to think about the involvement of the nucleosome.

### Half-nucleosome p53–ER grammar

p53 has been reported to bind preferentially near the nucleosome entry–exit region (*19*– *22*), suggesting that an ER motif approximately 73 bp away would potentially bind near the nucleosome dyad (Fig. 2A). To test whether this grammar could support a physically plausible complex, we modeled a nucleosome-bound p53–ER protein assembly using AlphaFold 3 (*23*). In the resulting model, ER occupied dyad-proximal DNA, whereas p53 was positioned near the nucleosome entry–exit region. This arrangement brought the two factors into close three-dimensional proximity, consistent with direct or indirect protein coupling across a half-nucleosome spacing (Fig. 2A) (*24*).

**Fig. 2.**
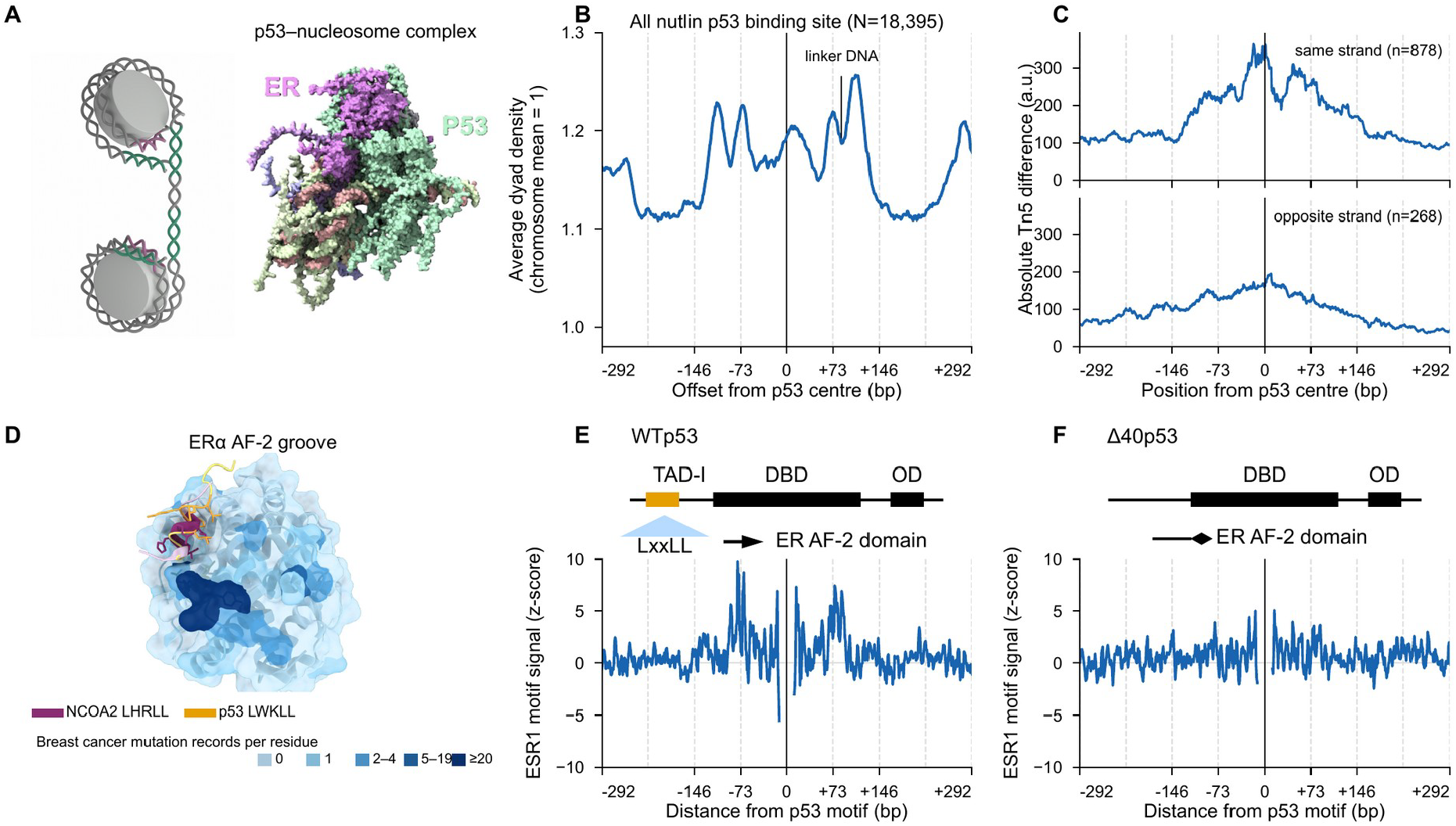
Structural and chromatin evidence for a 73-bp p53-ER grammar. **(A)** Schematic and AlphaFold 3 model of p53 and ER on a nucleosome, with ER dyad-proximal and p53 near the entry-exit region; ER is shown in purple and p53 in pale green in the structural model. **(B)** Mean MNase-seq dyad density around all nutlin-associated p53 binding sites in MCF7 cells (N = 18,395); the axis marks the p53 center and nucleosome-scale offsets. **(C)** Absolute nutlin-minus-DMSO Tn5-signal differences in MCF7 cells for same-strand (n = 878) and opposite-strand (n = 268) p53-ER configurations. **(D)** AlphaFold 3 model of p53 TAD-I at the ER-alpha AF-2 surface, overlaid with the NCOA2 peptide (PDB 1ZAF). The predicted p53 LWKLL motif (gold) overlaps the NCOA2 LHRLL-binding groove (purple); shading denotes somatic breast-cancer mutation records per residue. **(E and F)** ESR1 motif-signal profiles relative to p53 motifs in wild-type p53 (E) and Δ40p53 (F) ChIP-seq datasets. Curves show Nutlin − DMSO differences between separately aggregated profiles, independently z-scored using positions outside ±292 bp and displayed on shared axes. Input p53 ChIP-seq peak counts were 2,482 Nutlin and 699 DMSO peaks for wild-type p53, and 1,425 Nutlin and 659 DMSO peaks for Δ40p53.

To determine whether endogenous nucleosome distribution was consistent with this predicted grammar, average dyad density was mapped with MNase-seq in MCF7 cells and aggregated around all nutlin-associated p53 binding sites (n=18,395, Fig. 2B). The resulting profile showed phased dyad-density peaks flanking the p53 center at ±73 bp (and two additional peaks corresponding to exit-to-dyad distance plus a linker DNA length) and reduced dyad density near the central p53 binding site, consistent with preferential p53 positioning in linker or entry-exit DNA within an organized nucleosome context (Fig. 2B) (*24*).

To investigate whether p53 activation was associated with orientation-dependent accessibility changes, we aggregated ATAC-seq changes after nutlin-mediated p53 activation in MCF7 cells around p53 motifs and stratified the sites by ER motif orientation relative to p53. Same-strand p53–ER configurations (n=878) showed larger mean absolute accessibility changes than opposite-strand configurations (n=268). Differences were concentrated around the p53 motif and the predicted ER-positioned offset, supporting an association between the 73-bp configuration and regulated chromatin remodeling after p53 activation (Fig. 2C) (*25*).

### p53 TAD-I may interact with ER

The chromatin data suggested that p53 and ER complexes may cooperate within the same nucleosome. We therefore asked whether the two proteins contain a compatible interaction interface. Prior biochemical work has implicated the p53 N terminus in ER interaction, and this region contains the transactivation domain I (TAD-I) (*26, 27*). We modeled the p53 TAD-I peptide with the ER ligand-binding domain (LBD) using AlphaFold 3 and compared the predicted interface with experimentally characterized nuclear-receptor coactivator binding. The predicted p53 TAD-I interface overlapped the canonical ER AF-2 coactivator-binding surface and the position of a bound NCOA2 coactivator peptide (Fig. 2D). Because ER AF-2 recognizes LXXLL-like motifs (L, leucine; X, any amino acid), the LXXLL-like sequence in p53 TAD-I provides a potential molecular basis for this modeled interface (*23, 26, 28, 29*).

Mapping somatic breast cancer mutations across the ER LBD further highlighted this receptor-coupling region. Several recurrently altered residues occurred near the AF-2/coactivator surface, including structural neighborhoods relevant to endocrine resistance (Fig. 2D) (*30*). To test whether the p53 N terminus contributes to the 73-bp ER spacing grammar in chromatin, we compared ER motif-spacing profiles from wild-type p53 and Δ40p53 ChIP-seq. Δ40p53 is a p53 isoform that lacks the first 40 amino acids of p53, removing TAD-I and the LXXLL-like region. Full-length wild-type p53 showed a stronger flanking ESR1 motif signal near the nucleosome-scale registers than Δ40p53 when both profiles were independently standardized to distal positions and displayed on shared axes (Fig. 2, E and F). This attenuation in the Δ40p53 condition is consistent with a contribution of the p53 N terminus to the ER-associated nucleosome-scale grammar (*23, 29*).

### p53–ER grammar and cell cycle

CCND1, encoding cyclin D1, is a major ER target **(*2, 31*)**. In large breast cancer cohorts from TCGA and METABRIC **(*32, 33*)**, ER+ tumors consistently have significantly higher CCND1 expression, while p53 mutants show an ER-dependent decrease of *CCND1* expression (P=4.6e-04 for TCGA and P=0.003 for METABRIC, Fig. 3A), indicating that when ER is expressed, p53 could potentially cooperate with it on cyclin D regulation. Further, in the immortalized mammary epithelial cell line MCF10A, nutlin-induced p53 activation produces a 10-fold larger log-expression change with full-length wild-type p53 when compared to the Δ40p53 isoform (Fig. 3B), suggesting a role of the p53 N terminus in particular.

**Fig. 3.**
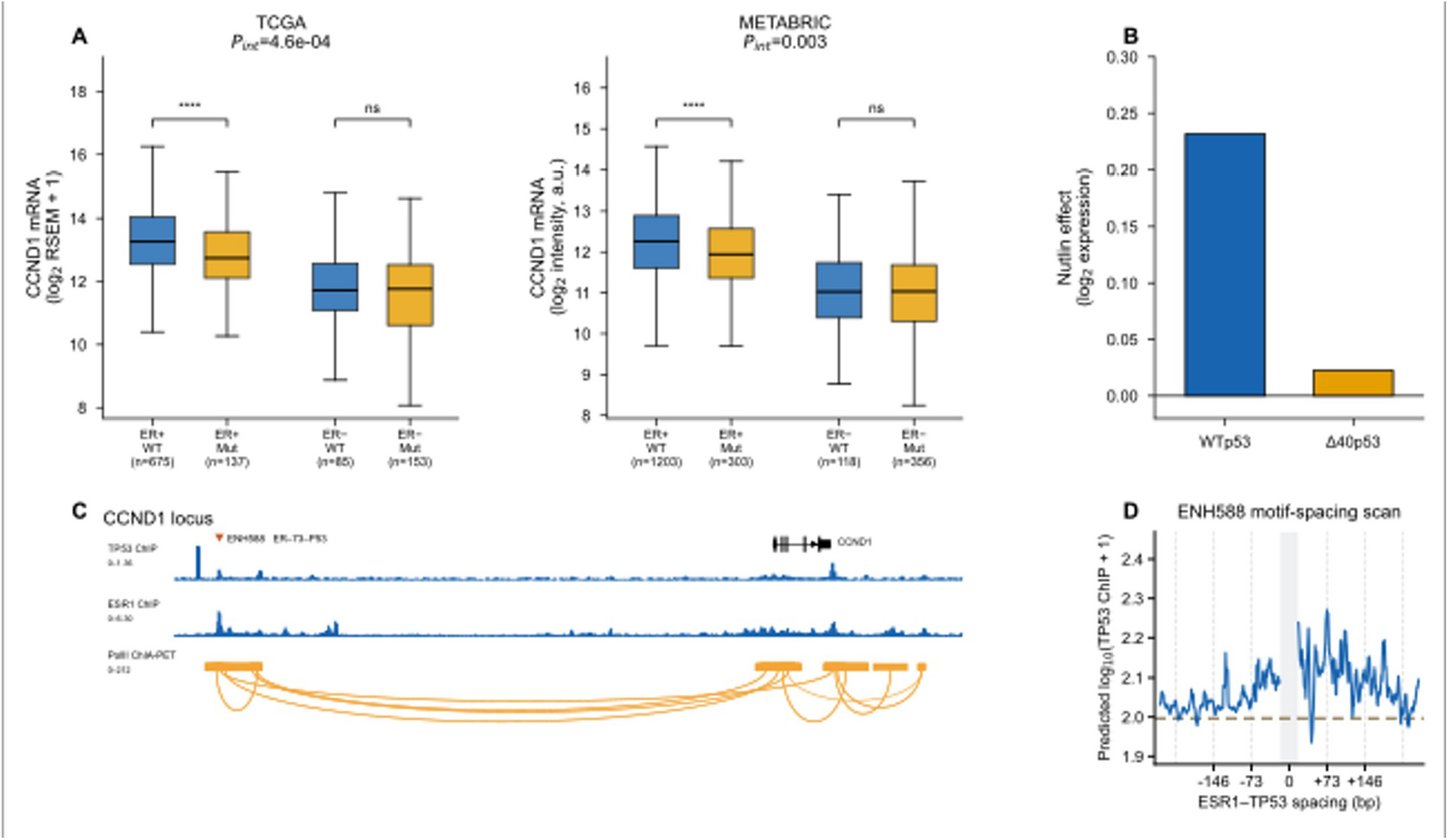
The p53-ER grammar links a CCND1 enhancer to cell-cycle states. **(A)** CCND1 expression by ER and TP53 status in TCGA (ER-positive: wild type, n = 675; mutant, n = 137; ER-negative: wild type, n = 85; mutant, n = 153) and METABRIC tumors (ER-positive: wild type, n = 1,203; mutant, n = 303; ER-negative: wild type, n = 118; mutant, n = 356). P_int_ values denote tests of the interaction between TP53 mutation status and ER status. **(B)** Nutlin-induced *CCND1* responses in wild-type-p53 and Δ40p53 MCF10A cells (n = 2 biological samples per genotype-treatment group). **(C)** TP53 and ESR1 ChIP-seq and Pol II ChIA-PET tracks at *CCND1*; ENH588 (triangle) contains the native ER-73-bp-p53 configuration. **(D)** ENH588 in silico spacing scan using TP53 ChIP-seq prediction model.

To study whether the 73-bp p53-ER grammar contributed to CCND1 regulation, we examined the entire gene locus. An ERα-bound enhancer, ENH588, was previously identified in a CRISPR screen whose disruption reduced ERα binding, CCND1 RNA and protein expression, estrogen-responsive CCND1 activation, and proliferation in ER-positive breast cancer cells (*2, 31, 34*). A 73-bp p53-ER grammar was found in this enhancer and TP53 and ESR1 occupancy were detected in breast cancer cells (Fig. 3C). An in silico spacing perturbation experiment (Methods) at this enhancer using our motif spacing-analysis deep learning model for TP53 ChIP signal showed that the native +73-bp p53-ER spacing leads to optimal p53 binding (Fig. 3D), suggesting a functional role for this grammar in this enhancer.

In tamoxifen-resistant p53-knockout cells, CCNE2 expression increased, with a significant treatment-by-TP53-genotype interaction (P = 0.00683; four-gene BH q = 0.0273; fig. S2A) and a similar trend for CCNE1. Separately, palbociclib-resistant T47D cells (mutant p53) showed a larger CCNE1 increase than resistant MCF7 cells (wild-type p53; fig. S2B). In patient samples, the ER-independent transcriptional subgroup increased from 5.0% before treatment to 20.7% after progression on combined endocrine and palbociclib therapy (*4, 29, 35*). These findings suggest cyclin E–CDK2 signaling as a potential route around ER–cyclin D–CDK4/6 dependence.

Hormone-receptor-driven cancers are enriched for wild-type TP53 at diagnosis and acquire TP53 loss as they progress to hormone independence. TP53 mutation is much less frequent in luminal A breast cancer (12%) but rises through luminal B and HER2-enriched to basal-like disease (∼80% TP53 mutations) (*32, 36*). A similar association between loss of hormone dependence and TP53 alteration is observed in prostate cancer, where TP53 alteration rises from 20% in localized castration-naïve disease to 37% in metastatic castration-naïve and 73% in metastatic castration-resistant prostate cancer (mCRPC) (*37, 38*). This pattern is consistent with a model in which p53-dependent chromatin opening supports nuclear receptor access to hormone response elements (*29, 39*), such that loss of p53 function is accompanied by loss of hormone receptor dependence.

### Genome-wide distribution of grammar by evolution of Alu sequences

Exploring the entire human genome for the recurrence of the p53-ER spacing grammar provides a clue to the evolution of this type of spacing. By stratifying Motiverse so as to plot annotated transposable element families (*40*), we found that Alu elements, derived from 7SL RNA approximately 60–65 million years ago and now represented by more than one million copies in the human genome, contributed strongly to receptor-motif enrichment around p53 (fig. S3A, **Supplementary Results**). To examine when these types of spacing arose, we compared grammar enrichment z scores with inferred TE expansion times across 17 primates and mouse, following the approach of Oomen et al. (*41*). Although the grammar was not exclusive to Alu family expansions, the AluS subfamily made the largest contribution to this primate grammar (Fig. 4A). These results identify Alu amplification as a mechanism for distributing recurrent p53–receptor motif grammar across primate genomes.

**Fig. 4.**
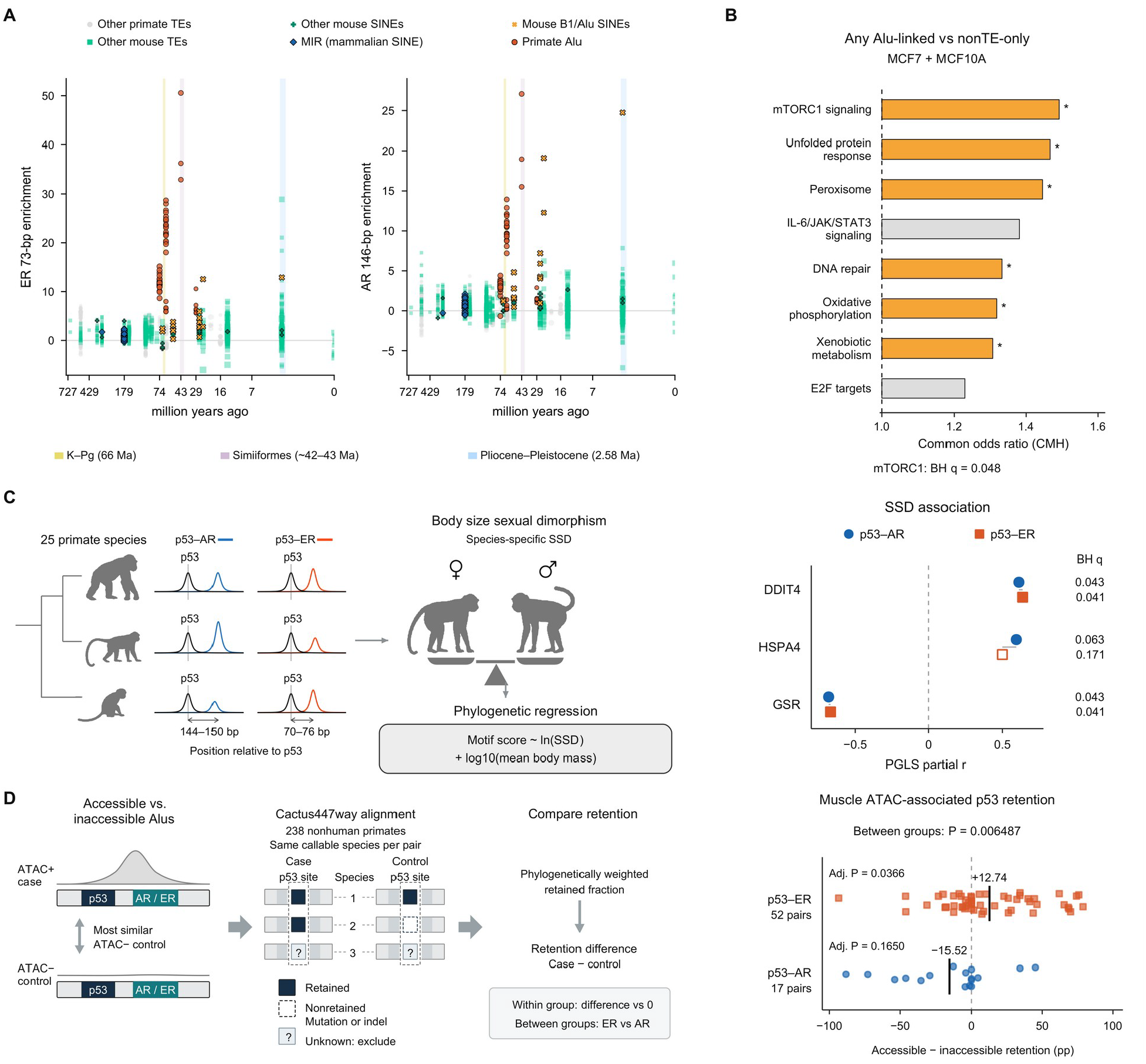
Transposable elements encode p53–receptor spacing grammar. **(A)** ER 73-bp and AR 146-bp motif enrichment across transposable-element families and inferred expansion times in 17 primates and mouse. Vertical bands mark the K–Pg boundary, Simiiformes divergence and the Pliocene–Pleistocene boundary. **(B)** Hallmark enrichment for genes within 100 kb of Alu-overlapping p53 peaks versus genes linked only to non-transposable-element p53 peaks in MCF7 and MCF10A cells. Bars show common odds ratios; the dashed line marks an odds ratio of 1. Asterisks indicate nominal P < 0.05; mTORC1 signaling passed multiple-testing correction (q = 0.048). **(C)** Schematic and body-mass-adjusted associations between p53–receptor motif scores and body size sexual dimorphism across 25 primates. Filled and open symbols indicate q < 0.10 and q ≥ 0.10, respectively. **(D)** Schematic and comparison of p53-site retention between muscle-accessible Alus and matched inaccessible controls. Each point represents a matched pair (52 ER and 17 AR pairs); black ticks mark means. Positive values indicate greater retention in accessible Alus. Statistical procedures are described in Methods.

Alu consensus comparisons placed the appearance of the grammar in AluS. We next asked whether Alu-associated p53 sites occurred near genes in particular pathways. Comparing high z-score genes within 100 kb of Alu-overlapping p53 peaks with inferred TE expansion times with those low z-scores p53 peaks not linked to the expansion times, identified mTORC1 signaling as the strongest hallmark enrichment (common OR = 1.49, BH q = 0.048; Fig. 4B), with the same direction of this association in human MCF7 and MCF10A cells. This is the case throughout primate evolution.

mTORC1 integrates nutrient and hormonal signals to regulate protein synthesis and muscle mass (*42, 43*). We therefore asked whether variation in p53–receptor motif scores near mTORC1-related genes was associated with body size sexual dimorphism, more commonly called sexual size dimorphism (SSD). Across 25 primate species, phylogenetic generalized least-squares regression accounted for average adult body mass and shared ancestry, as described by Rensch (*44*) (Fig. 4C, left). In the p53–AR screen, motif scores were positively associated with SSD at an Alu locus near DDIT4 (DNA damage induced transcript-4) (partial r = 0.612, q = 0.043) and negatively associated at a locus near GSR (glutathione-disulfide reductase) (partial r = −0.682, q = 0.043). HSPA4 (heat shock protein A4) showed a suggestive positive association (partial r = 0.595, q = 0.063; Fig. 4C, right). DDIT4 is a p53-regulated gene and encodes a stress-induced inhibitor of mTORC1 that reduces muscle metabolism under energetic stress (*45*). Additional locus analyses for these genes are presented in **Supplementary Results**.

We separately asked whether p53 motifs were more often retained across primates in Alu elements accessible in human muscle than in matched inaccessible Alus carrying the same nuclear receptor motif (Fig. 4D, left). Among Alus carrying a full estrogen-response element, retention was 12.74 percentage points higher in accessible elements (52 pairs; Holm-adjusted P = 0.0366). AR-motif-containing Alus showed a nonsignificant decrease of 15.52 percentage points (17 pairs; Holm-adjusted P = 0.165), with a difference of 28.27 percentage points between receptor classes (two-sided P = 0.00649; Fig. 4D, right). This pattern suggests different roles for ER and AR in muscle regulation.

### p53–receptor grammar in human muscle aging

Because hormone signaling, mTORC1 activity, and redox regulation each change during muscle aging (*46*), we examined p53–HR grammar in a single-cell transcriptomic and chromatin-accessibility atlas of 22 younger and older donors (*47*). In the original study, the atlas pointed to the same regulatory network predicted by GET in fetal skeletal myocytes (fig. S1C): AP1, FOXO1 and AR/GCR were among the principal regulators associated with muscle aging, while MDM2 expression was higher in older Type II myonuclei (fast-twitch skeletal muscle fibers, log_2_FC = +0.564; detected in 34.4% versus 16.4% of nuclei). This correspondence led us to search for p53–receptor grammar among the genes most closely associated with age.

Reanalysis of the single-cell RNA-seq data showed *MYLK4* (myosin light chain family member-4) as the strongest age-associated gene. This androgen-responsive regulator of muscle strength (*48*) showed the strongest relationship between myonuclear expression and donor age, with a marked decline in Type II myonuclei (Pearson r = −0.949, P = 1.90 × 10 ^−11^; Fig. 5A). The locus itself carried a matching chromatin signature, with an accessible promoter element at chr6:2,764,480–2,764,980 containing an exact 146-bp p53–AR grammar and overlapping TP53, ESR1 and AR ChIP-seq peaks. Its accessibility was lower in older Type II myonuclei (Δlog_2_(CPM + 1) = −0.730; Welch’s P = 0.0418; Fig. 5B). In mice, the p53 relevance for *Mylk4* expression was validated, where the expression is lower in satellite-cell-derived p53-null myoblasts than in wild-type cells (*49*) (Fig. 5C). *MYLK4* therefore connects the strongest age-associated muscle transcript in the atlas with the exact p53–AR grammar identified earlier in the study.

**Fig. 5.**
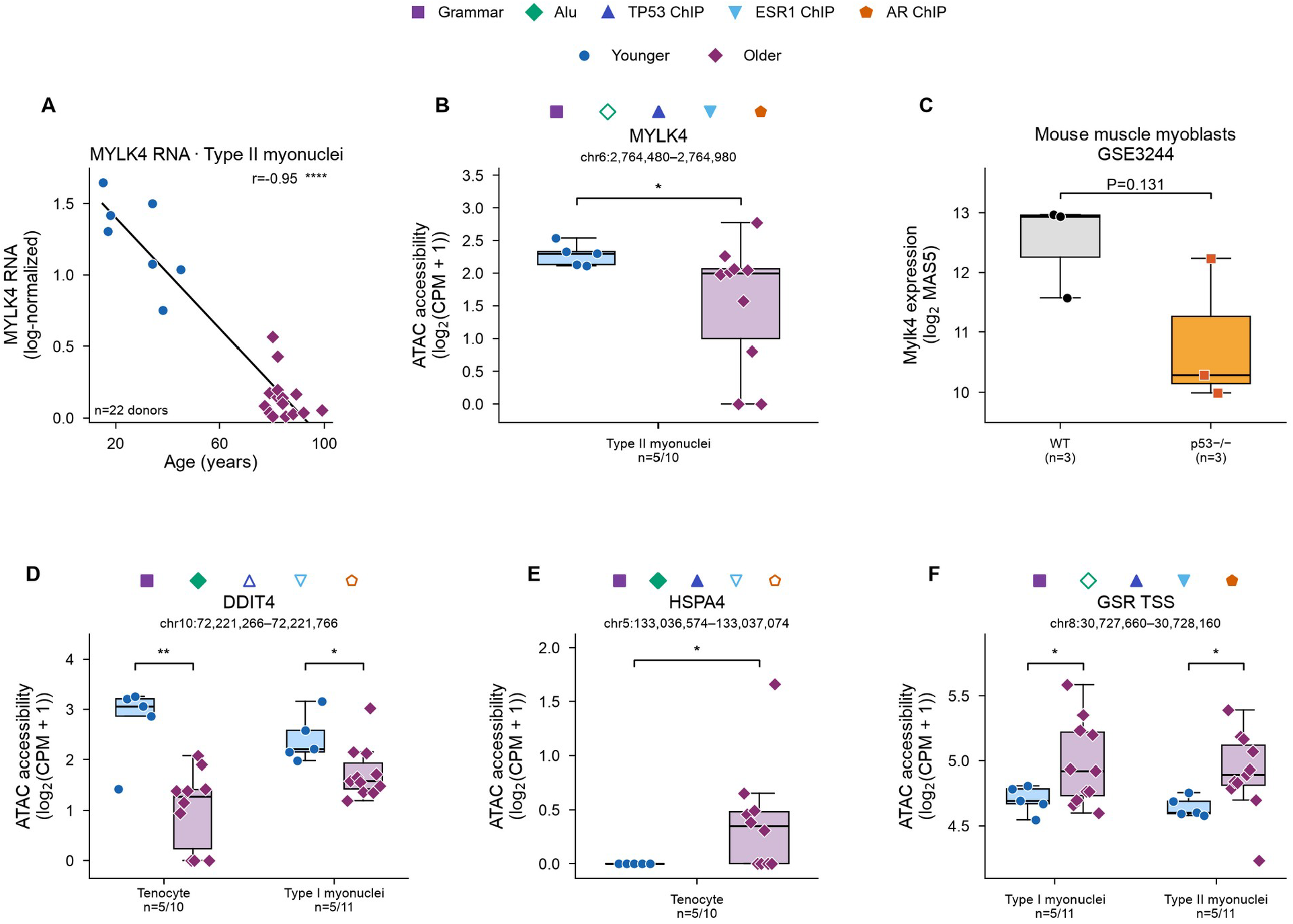
p53-receptor grammar and age-associated muscle regulation. **(A)** *MYLK4* RNA abundance in Type II myonuclei across donor age; points denote younger (n = 7) and older (n = 15) donors, with the fitted correlation shown. **(B)** ATAC-seq accessibility at the *MYLK4* promoter interval chr6:2,764,480-2,764,980 in younger (n = 5) and older (n = 10) Type II myonuclei; symbols above the locus indicate the 146-bp grammar, Alu overlap, and TP53, ESR1 and AR ChIP-seq support. **(C)** Mylk4 expression in wild-type and p53-null mouse muscle myoblasts (GSE3244). **(D–F)** Age-group ATAC-seq accessibility at *DDIT4* in tenocytes and Type I myonuclei (D), and *HSPA4* in tenocytes (E), and the GSR transcription start site in Type I and Type II myonuclei (F). Due to filtering on cell count (at least 20 nuclei), ATAC-seq donor counts were 5 younger and 10 older for HSPA4 tenocytes and DDIT4 tenocytes, and 5 younger and 11 older for DDIT4 Type I and GSR Type I and Type II myonuclei.

We then returned to *DDIT4, HSPA4* and *GSR*, the mTORC1-related genes nominated by the primate SSD analysis. Each of them contained an age-responsive accessible element supported by Alu overlap or p53–receptor grammar. *DDIT4* showed the largest accessibility shift. The peak (chr10:72,221,266–72,221,766) overlaps a p53–AR Alu. Accessibility was markedly lower in older tenocytes (Δ = −1.73; P = 0.0031) and was also reduced in older Type I myonuclei (slow-twitch skeletal muscle fibers, Δ = −0.67; P = 0.0318; Fig. 5D). At the HSPA4 locus, an age-responsive peak (chr5:133,036,574–133,037,074) overlaps two Alu elements that carry p53–AR and p53–ER grammars. This interval also overlaps TP53 ChIP-seq peaks from 82 ChIP-Atlas experiments from various cell types. Its accessibility increased in older tenocytes (Δ = +0.397; P = 0.0354; Fig. 5E), extending the association to the aging tendon compartment. *The GSR* promoter (chr8:30,727,660–30,728,160) contains a p53–ER motif pair separated by 70 bp, and overlaps TP53, ESR1 and AR ChIP-seq peaks. Its accessibility increased with age in both Type I (Δ = +0.274; P = 0.0245) and Type II myonuclei (Δ = +0.267; P = 0.0179; Fig. 5F). Together, these results identified age-responsive p53–receptor elements at *MYLK4* and at the SSD-associated *DDIT4, HSPA4* and *GSR* loci across human myonuclei and tenocytes.

## Discussion

In this work, we identified a nucleosome-scale regulatory grammar that organizes cooperation between p53 and steroid hormone receptors. Distinct p53–ER and p53–AR motif grammars recur at approximately one-half and one nucleosome, respectively, linking chromatin structure to transcription-factor cooperation, cell-type-specific regulation, and evolutionary innovation. We further showed that these motif grammars were disseminated by transposable elements, providing a mechanism by which regulatory grammars can emerge and expand during genome evolution. The function of full-nucleosome spacing between p53 and AR calls for further study. Beyond p53–ER and p53–AR interactions, the conserved 37-bp spacing between p53 and KMT2A motifs in both Alu and LINE1 elements (fig. S3) suggests that additional nucleosome-dependent grammars remain to be identified and comprehended through the same strategy.

Why p53 evolved as a central partner for steroid hormone receptors remains an open question. The deep evolutionary conservation of the p53 family, together with its roles in stress surveillance, chromatin remodeling, and stem-cell biology, places it at the intersection of hormone signaling, metabolism, and tissue homeostasis. Consistent with this broader metabolic role, individuals with Li-Fraumeni syndrome carrying germline TP53 mutations exhibit increased skeletal-muscle oxidative phosphorylation, supported by patient-derived cells and a mouse model (*50*), which is linked to increased glutathione levels, mTOR–Rheb association, and improved muscle function and recovery (*51*). Interestingly, GSR directly participates in glutathione metabolism and exhibits sex-dependent regulation (*52*). Perturbation of this p53–receptor regulatory hub in Li–Fraumeni syndrome may preferentially predispose patients to ER-positive breast cancer, consistent with an important role for p53–ER cooperation in maintaining mammary epithelial identity. One intriguing possibility is that reduced p53 dosage destabilizes cooperative regulatory states established by these grammars, thereby lowering the robustness of differentiated luminal programs giving rise to ER+ breast cancers at a higher rate (90% of women) and a younger age (20-40 years old) than women with inherited diploid TP53 genes.

More generally, our findings suggest that transcription factor cooperativity is encoded not only by compatible protein interfaces or individual binding motifs, but also by their higher-order spatial organization in DNA. After a new protein interaction is established, constructing a large number of compatible binding sites through individual mutations is probabilistically prohibitive. In this view, transposable-element expansion provides an evolutionary process that establishes recurrent regulatory architectures, whereas chromatin remodeling dynamically selects which grammars are accessible in a given cell state based on expressed transcription factors. In this case the evolutionary selection pressures may well have selected for some types of sexual dimorphism. Indeed the p53–AR–mTORC1 interaction demonstrating loss of muscle functions with aging in primates permits younger male primates to take over mating ensuring higher quality of spermatogenesis is passed on in the species. The distributed organization of regulatory grammars across repetitive elements resembles attractor-based models of gene-regulatory networks and associative-memory models, first described by Hopfield (*53*) and expanded by Krotov and Hopfield (*54*), in which stable cellular identities emerge from cooperative interactions among many regulatory nodes. Although speculative, such a framework offers a mechanistic perspective for how regulatory grammars could contribute to the establishment and maintenance of differentiated cellular identities (see Supplementary Discussion).

Finally, our study provides a general methodological framework for discovering cooperative regulatory grammars that can be applied to other cellular contexts and TFs. This is particularly useful for comparative regulatory genomics because noncoding sequences can diverge substantially at the base level while preserving functional motif relationships, whereas transcription-factor binding does not require an exact match to a single consensus sequence. Motiverse is therefore broadly applicable across diverse transcription-factor families and may be extended to less-studied species in which DNA-binding domains and motif preferences remain evolutionarily conserved. Together with the accompanying comparative transposable element atlas spanning 96 species (https://fuxialexander.github.io/motiverse/atlas/), these resources provide a general framework for discovering and comparing cis-regulatory grammars across genomes and throughout evolution.

## Use of generative AI tools

OpenAI Codex was used to assist with language editing and data analysis. The authors initiated, reviewed and verified all scientific content, analyses, interpretations and citations.

## Data availability

Public functional-genomics datasets used in the current figures and analyses are GSE111009 (MCF10A Nutlin/DMSO TP53 ChIP–seq for pathway enrichment), GSE147700 (MCF10A p53 ChIP–seq), GSE147701 (MCF10A RNA-seq), GSE250017 (MCF7 nutlin ATAC-seq), GSE282874 with GSM8651880, GSM8651881 and SRR31516404 (MCF7 MNase), GSE292363 (MCF7 p53/tamoxifen RNA), GSE222367/SRP416348/PRJNA920517 (palbociclib resistance), and GSE86164 with GSM2296275/SRX2060922 (MCF7 nutlin p53 ChIP–seq used for the Motiverse MCF7 profiles), OMIX004308-05 and OMIX004305-05 (Human Muscle Aging Atlas). TCGA breast cancer (brca_tcga) and METABRIC (brca_metabric) expression, TP53 mutation and *CCND1* copy-number data were accessed through cBioPortal. Motifs were obtained from HOCOMOCO v14; repeat annotations and sequences were obtained from RepeatMasker/UCSC for the 17 assemblies listed in **Methods**; the primary primate phylogeny was obtained from the Mammal Phylogeny Project/Upham resource. Key coordinate systems are hg38 unless stated otherwise; ENH588 is chr11:69,514,361–69,516,409 with TP53 chr11:69,515,374–69,515,395 and ESR1 chr11:69,515,450–69,515,465.

## Code availability

Motiverse is available at https://github.com/fuxialexander/motiverse; GET code and the cell-state catalogue are at https://github.com/GET-Foundation/gcell. The sequence-model configuration and checkpoint are deposited in the Motiverse repository.

## Author Contributions

X.F., A.L. and R.R. conceived the study and designed the analyses. X.F. developed Motiverse and performed computational and biological analysis. Q.C. trained the TP53 ChIP-seq model with advice from X.F. X.F., A.L., and R.R. wrote the manuscript with helpful suggestions from Q.C. and P.S.

