## Supplementary Materials for "Nucleosome-scale p53–hormone receptor grammar distributed by Alu elements"

#### Materials and Methods

##### *Motiverse motif-spacing analysis*

Regulatory intervals were scanned with aligned HOCOMOCO v14 position-weight matrices, including reverse complements. In the one-to-all mode, a query P53.H13CORE.0.P.B call at a mapped $P < 10^{-4}$ threshold anchored a $\pm 500$-bp window. Convolutional motif scores were filtered using the corresponding HOCOMOCO P-value mapping; aligned-kernel padding was corrected by each motif core center. For every anchor, Motiverse retained partner-motif identity, score, strand and signed center-to-center distance, producing a motif × 1,001-position array (or subset × motif × position array). The displayed aggregate profiles are three-base-pair-smoothed means standardized to the distal positions $|\text{distance}| > 292$ bp.

##### *ChIP-seq Model Training*

TP53 ChIP-seq data for MCF7 cells treated with nutlin were obtained from GSE86164. The training target was generated from the nutlin-treated TP53 ChIP profile after input subtraction and cPeaks normalization (details described in **Chromnitron** (55)): $\text{target} = \max((\text{TP53 ChIP} - 1.4522 \times \text{input}) / 27.3193, 0)$, where the input scaling factor was estimated from background genomic regions as the ratio of mean raw TP53 ChIP signal to mean raw input signal, and the cPeaks-derived scale factor was estimated from the highest signals in cPeaks regions. No library-size normalization was applied.

Peaks were called from the processed profile, merged, and recentered to 2048 bp windows by IQR-based detection. Peak regions were retained if they had $\text{SumValue} \geq 50$ and $\text{MaxValue} \geq 0.2$. Background regions were sampled from the complement of cPeaks at 30% of the retained peak count. The final dataset contained 306,887 regions, including 236,067 peak regions and 70,820 background regions. Chromosome 15 was held out for validation, and all remaining autosomes plus chromosome X were used for training.

The model consisted of a sequence-only convolutional neural network initialized with 3,222 aligned motif kernels from HoCoMoCo v13. Motif convolution outputs were passed through group normalization and a stack of dilated residual convolutional blocks with kernel size = [1, 2, 3, 5, 11, 41, 81, 167]. The shared sequence representation was used by two output heads: a nucleotide-resolution profile head and a scalar count head. Predictions were evaluated over the central 1000 bp of each input window. The training objective combined a BPNet-style profile loss with a count loss. For the profile loss, the observed cPeaks-normalized TP53 signal was smoothed with a 20 bp window, normalized to a probability distribution across the central 1000 bp, and compared with predicted profile logits using cross-entropy. Profile loss was applied only to regions with $\text{SumValue} \geq 30$. The count loss was mean squared error between predicted and observed $\log_{10}(\text{signal sum} + 1)$ and was applied to all valid regions. The total loss is defined as: profile loss + $10 \times$ count loss.

The model used only DNA sequence as input. For each region, a 2048 bp one-hot encoded hg38 sequence window was extracted. During training, random positional augmentation was applied by shifting the window center by up to $\pm 500$ bp, and reverse-complement augmentation was applied independently.

Models were trained with AdamW using a learning rate of $1 \times 10^{-4}$, batch size 32 and 32-bit precision. Training was run for up to 100 epochs with early stopping on validation loss using a patience of 20 epochs.

#### *Spacing perturbation analysis*

The ENH588 locus spacing perturbation scan used the sequence at hg38 chr11:69,514,361–69,516,409. The fixed TP53 motif was chr11:69,515,374–69,515,395 and the native ESR1 motif was chr11:69,515,450–69,515,465, a +73-bp center-to-center configuration. The native ESR1 sequence was removed, the flanks were stitched, and the same motif was reinserted at each allowed offset; motif-overlap offsets were recorded as missing. The deletion baseline is the stitched ESR1-deleted sequence. Each value is a five-base-pair centered rolling mean of $\log_{10}(\text{TP53 ChIP} + 1)$.

#### *GET context prioritization*

Cell-state regulatory hypotheses were derived from GET transcription-factor interaction matrices. The full retained catalogue contained 4,440 p53-family-to-receptor rows spanning 185 cell identifiers or cell types. P53-like motif clusters (P53-like/1, /2 and /3) were queried against ER-compatible clusters (NR/7, NR/12 and NR/17) and an AR-proxy cluster (NR/20). For Figure 1, each signed weight was standardized within its cell state against all off-diagonal edges and ranked by its absolute z score; receptor-expression predictions were displayed separately. Mammary luminal and skeletal-muscle-related states were inspected after this catalogue-wide ranking.

#### *Structural and mutation context*

AlphaFold 3 models were used as hypothesis-generating structural predictions of the p53–ER–nucleosome complex and the p53 TAD-I–ERα ligand-binding-domain interface. ChimeraX was used to visualize the predicted structures.

#### *MNase and ATAC analyses*

MCF7 MNase dyad profiles were obtained from GEO GSE282874. Processed dyad BigWigs GSM8651880 and GSM8651881 and retained reprocessed FASTQs including SRR31516404 were used. Signal was aligned on p53 motif centers. Strict ER73 anchors required an ER-like center within $\pm 3$ bp of either 73-bp offset; off-phase controls contained qualifying ER-like motifs outside this class. Half-wrap, p53-center and outer-dyad windows were compared with far-background windows. Group contrasts used two-sided Mann–Whitney tests; co-occurrence used upper-quartile signal classes, Fisher exact tests and rank or linear correlations as labeled.

MCF7 ATAC-seq from GSE250017 was compared between DMSO and nutlin-3a. Tn5 insertion profiles were centered on p53 or ER-like motifs and stratified by partner presence, signed center-to-center distance and same-versus-opposite strand orientation. Figure 2 reports the absolute nutlin-minus-DMSO insertion difference around the p53 center and predicted ER73 positions. Since MNase and ATAC profiles measure aggregate chromatin signals rather than individual molecules, the observed profiles could result from bulk aggregation.

For the wild-type p53 and $\Delta 40$p53 comparison (Fig. 2, E and F), ESR1 motif profiles were aggregated separately for Nutlin and DMSO p53 ChIP-seq peak sets from MCF10A cells (GSE147700), and the DMSO profile was subtracted from the Nutlin profile within each genotype.

The inputs comprised 2,482 Nutlin and 699 DMSO unique peak intervals for wild-type p53, and 1,425 Nutlin and 659 DMSO intervals for $\Delta 40p53$.

#### *Repeat and cross-species analyses*

RepeatMasker/UCSC rmsk annotations were paired with 17 assemblies: calJac4, chlSab2, gorGor6, hg38, macFas5, micMur2, nasLar1, nomLeu3, otoGar3, panPan3, panTro6, papAnu4, papHam1, ponAbe3, rheMac10, rhiRox1 and saiBol1. Copies were grouped by RepeatMasker family/class and oriented by the annotated strand; truncated copies were excluded.

For human Alu tests, 876,431 full-length-like copies were scanned for p53, ER-like and AR-like hits. Cross-species examples used mm10 B1\_Mm (n = 1,622) and dm6 IDEFIX\_I-int (n = 1,325).

#### *Expression and tumor analyses*

MCF10A expression analysis used GSE147701, with two biological samples in each genotype-by-treatment group. Welch contrasts between such small groups were treated as exploratory. For the MCF7 analysis, rlog expression from GSE292363 was analysed separately for each gene by ordinary least squares:

$$\text{expression} \sim \text{genotype} + \text{tamoxifen} + \text{genotype} \times \text{tamoxifen}.$$

The analysis included four untreated and five tamoxifen-treated wild-type samples, and four untreated and four tamoxifen-treated p53-knockout samples. Two-sided coefficient P values were adjusted by the Benjamini–Hochberg method across the four displayed genes.

For the palbociclib-resistance analysis, processed counts from GSE222367 (BioProject PRJNA920517; SRA study SRP416348) were converted to $\log_2(\text{CPM} + 0.5)$. Parental cells were compared with independently selected, long-term palbociclib-resistant MCF7 and T47D states. Each cell state and drug concentration included three biological samples.

ER-positive primary breast tumors were obtained through cBioPortal from brca\_tcga and brca\_metabric. TP53-mutant samples carried any reported nonsynonymous TP53 mutation. Expression profiles were brca\_tcga\_rna\_seq\_v2\_mrna ($\log_2[\text{RSEM} + 1]$) and brca\_metabric\_mrna (normalized expression); CCND1 copy number came from the corresponding cBioPortal study. Unadjusted TP53-group comparisons used two-sided Mann–Whitney tests. Sensitivity models used ordinary least squares with TP53 status and CCND1 copy number, plus molecular-subtype indicator variables for METABRIC. Sample sizes were TCGA 675 wild type and 137 mutant and METABRIC 1,203 wild type and 303 mutant (30, 56).

#### *Pathway enrichment*

We restricted the analysis to two mammary-cell studies with endogenous, unengineered TP53: MCF7 cells from GSE86164 and MCF10A cells from GSE111009. Within each study, ChIP-Atlas TP53 peaks detected after Nutlin treatment but absent from the matched DMSO control were retained at a score threshold of $\geq 100$ and linked to genes within 100 kb. We compared all genes linked to Alu-overlapping TP53 peaks, including genes also linked to non-TE peaks, with genes linked only to non-TE TP53 peaks using a one-sided Cochran–Mantel–Haenszel analysis, with false-discovery rates calculated across all 50 Hallmark gene sets using the Benjamini–Hochberg method.

#### *Phylogenetic regression*

At each locus, the p53 score was evaluated at the human-register homologous position. AR and ESR1 components were the strongest motif scores 144–150 bp and 70–76 bp, respectively, from p53 on either side; component scores and their sums were retained as separate endpoints. Sexual-size dimorphism (SSD) was the male/female adult body-mass ratio, entered into the model as $\ln(\text{SSD})$, and mean adult mass was $\log_{10}$-transformed. Analyses used 25 included primate species after collapsing duplicate *Hylobates* labels and excluding *Papio kindae* because required covariates were unavailable. Shared ancestry used the Mammal Phylogeny Project/Upham node-dated maximum-clade-credibility tree under Brownian covariance, with the fossilized-birth-death tree as a sensitivity analysis.

For each receptor family, the reported screen comprised 24 loci, with motif score regressed on SSD and $\log_{10}$ mean adult mass included as a covariate. Empirical support used 100,000 phylogeny-aware shuffled datasets that preserved the body-mass relationship. P values were Benjamini–Hochberg-adjusted separately across the 24 loci in each receptor screen.

#### *Muscle-accessible Alu motif retention*

Accessible and inaccessible Alus were paired within receptor motif class using the original nearest-control matching design. Greedy matching selected the closest unused eligible control, processing cases with fewer candidates first. Retention differences were calculated over each pair's shared callable nonhuman primates (minimum ten), using fair-proportion phylogenetic weights. Within-group tests used two-sided paired swaps, with Holm correction across the three-group exploratory family. The full-ERE-minus-AR contrast used a two-sided wild cluster bootstrap with 999,999 replicates and 61 shared-Alu components.

#### *Human muscle aging and candidate-locus analysis*

Processed snRNA-seq and snATAC-seq data from the Human Lifespan Muscle Atlas were obtained from OMIX004308-05 and OMIX004305-05. For MYLK4 RNA, log-normalized expression was averaged across Type II myonuclei within each donor and correlated with continuous donor age using a two-sided Pearson test. For ATAC, union-peak counts were summed within each donor and cell type, normalized to the corresponding PeakMatrix library size, and expressed as $\log_2(\text{CPM} + 1)$. Donor–cell-type combinations containing at least 20 nuclei were included, with a minimum of three donors in each age group. The atlas Adult and Old groups are labelled Younger and Older in the figure. Group differences were evaluated using two-sided Welch's t-tests, with donors as biological replicates. The GSR promoter p53 motif was scanned at $P \leq 0.005$. ChIP support was defined by factor-matched interval overlap with the official hg38 ChIP-Atlas all-cell assembled tracks at $q < 10^{-5}$. Mouse *Mylk4* expression was obtained from GSE3244/GPL83 and compared between three wild-type and three p53-null satellite-cell-derived myoblast samples using a two-sided Welch's t-test.

#### *Statistical analysis*

All tests were two-sided unless an enrichment alternative was explicitly specified. Fisher exact tests were used for motif-register and pathway contingency tables; two-sided Mann–Whitney tests for independent signal or tumor-expression groups; Spearman correlation for monotonic profile associations; Pearson correlation only for prespecified approximately linear continuous comparisons; and ordinary least squares for the factorial RNA and tumor sensitivity models.

described above. Bootstrap intervals resampled independent motif anchors. Benjamini–Hochberg $q$ values were calculated for multiple-hypothesis testing where applicable.

### **Supplementary Results**

#### ***Repeat-family patterns and inferred expansion times***

TE-stratified Motiverse profiles showed weaker but related enrichment in LINE-1 elements alongside the stronger Alu patterns (Fig. S3A). The inferred timing of Alu-associated grammar expansion fell near the K–Pg boundary (approximately 66 million years ago) and Simiiformes divergence (approximately 43 million years ago; Fig. 4A). Mouse SINEs and other TEs showed later inferred expansions, around 29 million years ago, while precursor sequences contained related motif grammar.

#### ***Phylogenetic regression locus analyses***

The p53–AR component analysis at DDIT4 suggested a positive AR contribution ($q = 0.052$), whereas the positive HSPA4 composite-score association was driven by variation in its p53 component. The p53–ER screen also identified the Alu near DDIT4 at hg38 chr10:72,180,028–72,180,230 ($q = 0.041$; Fig. 4C). A separately examined p53/AR-bound distal enhancer near DDIT4 (hg38 chr10:72,297,273–72,299,152) showed a positive p53–AR score association with SSD (partial $r = 0.485$, $q = 0.079$). This interval interacts with the DDIT4 promoter in MCF7 Pol II ChIA–PET and has active enhancer marks and p53/AR occupancy annotations across tissues, including skeletal muscle. Genome alignment across 447 species shows that the enhancer is conserved across placental mammals.

#### ***Additional repeat-associated grammars***

Alu-centered patterns were not limited to ER and AR. KMT2A and JUN also showed structured subnucleosomal enrichment around p53 (Fig. S3A), suggesting that repetitive elements may encode additional spatial relationships among regulators. Repeat and motif annotations used RepeatMasker and HOCOMOCO (40, 57).

Tracing Alu consensus sequences from 7SL RNA through the free left Alu monomer (FLAM) indicated that the 7SL-derived sequence lacked a strong p53 site, whereas a p53-like motif appeared in FLAM and the strongest modern p53 spacing emerged near position 150 in AluS. The corresponding ER motif appeared in AluS and was absent from older AluJ and FLAM references. The AR motif near position 296 appeared to derive from poly(A)-proximal AluS sequence and was not evident in AluJ, FLAM or 7SL-derived DNA. These observations support acquisition or retention of composite grammar after the ancestral Alu scaffold emerged.

Furthermore, a systematic screen in mouse and fly identified mouse B1\_Mm elements and *Drosophila* IDEFIX\_I-int elements with 73-bp ER-like motif enrichment (1,622 and 1,325 p53 anchors, respectively; Fig. S3B). These observations indicate that related p53-centered spacing relationships can arise in distinct repeat families outside primate Alu elements.

### Supplementary Discussion

When a protein complex composed of two transcription factors binds to two different response element (RE) sequences in DNA, this can give rise to avidity and increased binding stability (58). This resembles a bistable switch that can encode one bit of memory. Hopfield associative memory (53), by contrast, acts at the network level and can encode many bits of memory. Here, such memory could arise from distributed, coupled nodes of Alu sequences. Binding avidity together with network attractor capacity could produce a differentiated state, or attractor, that is stable over time. Loss of this state, through TP53 mutation for example, could permit transition to a new proliferative state, contributing to ER+ cancers (29, 35). In this framework, cellular memory regulates and restrains the cell cycle, whereas loss of memory promotes proliferation. Thus, transcription factor complexes together with distributed repetitive response elements positioned at defined intervals could generate forms of associative memory (53) and dense associative memory (54) analogous to those described mathematically for neural networks.

In living systems, this type of memory could operate over two timescales. First, evolutionary sequence rewiring of repetitive elements occurs slowly; once fixed, these sequences can subsequently be altered by mutation and acted upon by selection (9). Second, epigenetic chromatin modifications can act rapidly to alter chromatin accessibility for transcription factor binding and are reversible through regulated, enzyme-catalyzed chemical reactions (29). The grammar and architecture of the transcriptional programs described in this manuscript may therefore be related to processes that resemble the associative memory first described by Hopfield (53) and later extended by Krotov and Hopfield (54) in mathematical models of neural networks that contributed to the development of modern machine learning. One possibility is that this form of cellular memory contributes to observations such as the lower incidence of ER+ breast cancer among women who undergo childbirth and breastfeeding at younger ages compared with nulliparous women, potentially because pregnancy and lactation promote a stable differentiated state in mammary ductal cells (59). In Li-Fraumeni syndrome (LFS), individuals carry one germline mutant TP53 allele and therefore retain only one wild-type TP53 allele. ER+ breast cancer is the most common cancer observed in affected women, with very high lifetime penetrance and frequent onset between 20 and 40 years of age; loss or functional inactivation of the remaining wild-type TP53 allele can contribute to tumor development and may underlie some of the earliest cases of breast cancer in women with LFS; after cancer develops in one breast, the risk of a second primary breast cancer is also substantial. In this setting, the protective association with parity appears to be reduced, and the benefit associated with breastfeeding may also be partially compromised. Evidence that functional p53 dosage contributes to these effects is based on one human study together with mouse studies and is not yet sufficiently strong to be clinically actionable (50, 59) but it nevertheless further links p53 biology to ER+ breast cancer.

The association between wild-type p53 function, sexually dimorphic transcriptional programs, and susceptibility to ER+ breast cancer raises an evolutionary possibility. Regulatory interactions involving p53 and estrogen receptor may have been retained because they confer physiological or reproductive benefits earlier in life, while creating a vulnerability to ER+ breast cancer later in life—an example of antagonistic pleiotropy. The experimental results presented here are consistent with this possibility, but establishing such an evolutionary mechanism will require additional testing.

The earliest p53-family transcription factor identified on the basis of DNA-sequence homology is found in modern-day choanoflagellates and cnidarians, lineages that diverged near the emergence of animal multicellularity approximately 800 million years ago (1). In *Drosophila*, the single p53-like gene displays sexually dimorphic functions during development, with distinct effects in males and females (60). Sexually dimorphic regulation involving p53-associated pathways continues to be observed in mammals, including humans (29, 35, 61). In primates,

TP53 participates broadly in stress surveillance, including in muscle and hematopoietic stem cells, where hormone signaling, metabolism, immune function, and sex-biased physiology intersect. In humans, p53 regulates genes encoding epigenetic and chromatin-modifying activities that can alter accessibility of estrogen and androgen response elements located within the same nucleosome, including sites approximately 73 bp and 146 bp from a p53-binding site. In addition, the TP53 gene contains a bone-marrow-associated transcriptional enhancer in its fourth intron (61) , potentially linking TP53 regulation to sexually dimorphic features of the immune system.

### Supplementary Methods

#### *Motiverse algorithm details*

Motiverse separates anchor selection from partner profiling. First, sequence windows are fetched for input intervals in a declared genome assembly. Second, aligned HOCOMOCO kernels and their reverse complements are convolved with each window. Third, a qualifying query-motif hit is chosen as the anchor and its corrected motif-core center is set to position zero. Fourth, every partner hit is written at its signed center-to-center offset with motif identity, strand and score. Fifth, anchor-level arrays are reduced within declared biological subsets; smoothing, distal-background standardization and register-specific tests are performed only after this anchor-level representation has been saved.

Conceptually, the procedure pseudocode is:

```
for interval in intervals:
    sequence = fetch(genome, interval ± window)
    scores = conv1d(sequence, aligned_motifs_and_RC)
    anchor = qualifying_query_hit(scores[query])
    for motif in motifs:
        offset = corrected_center(motif_hit) -
        corrected_center(anchor)
        profile[motif, offset] += score
        counts[offset] += 1
```

The reported profile is profile/counts. For display in the manuscript, profiles were smoothed over three base pairs and standardized against positions more than 292 bp from the anchor. Both strands were combined unless a panel was explicitly strand-specific.

#### *Runtime benchmark*

On one NVIDIA GeForce RTX 5090, a measured fresh p53-all run took 195.14 s (3.25 min) to identify and save p53 anchor positions and then generate the downstream profiles for 3,222 motifs across 100 subsets. The anchor-identification/cache-construction component took 58.44 s. Repeating the same analysis while reusing those saved p53 anchor positions took 92.44 s; partner-motif scoring and the 1,001-position output curves were still recomputed. For comparison only, two older saved p53-ER/FOXA1 logs imply 4.1–5.3 min for observed plus circularly permuted scanning of 30,294 peaks.

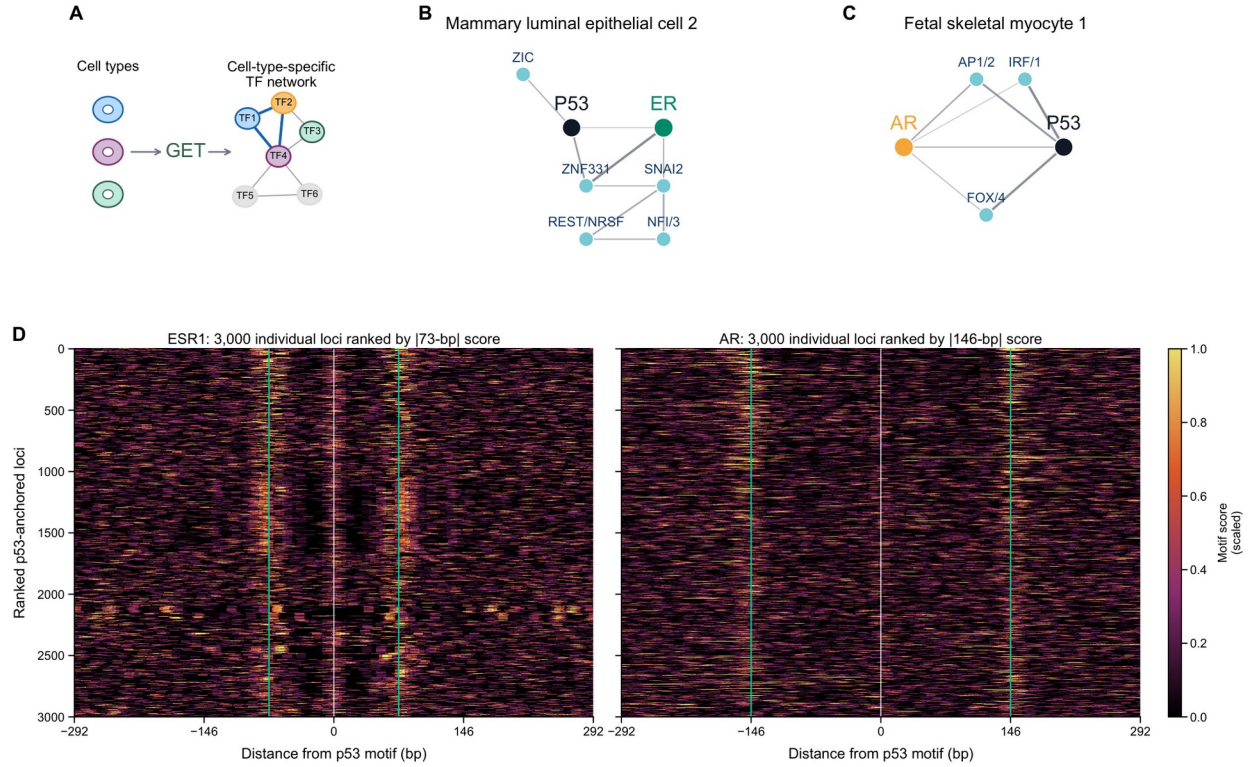

**Fig. S1. Cell-context dependence of p53-associated receptor grammar. (A)** GET workflow for cell-type-specific transcription-factor networks. **(B and C)** GET core regulation network for mammary luminal epithelial cell 2 (B) and fetal skeletal myocyte 1 (C). **(D)** Per-locus ESR1 and AR motif-score heatmaps around p53 in MCF7 nutlin-treated TP53 ChIP-seq peaks. The 3,000 regions with the largest 73-bp or 146-bp motif scores were visualized. White lines mark p53; teal lines mark +/-73 bp for ESR1 and +/-146 bp for AR.

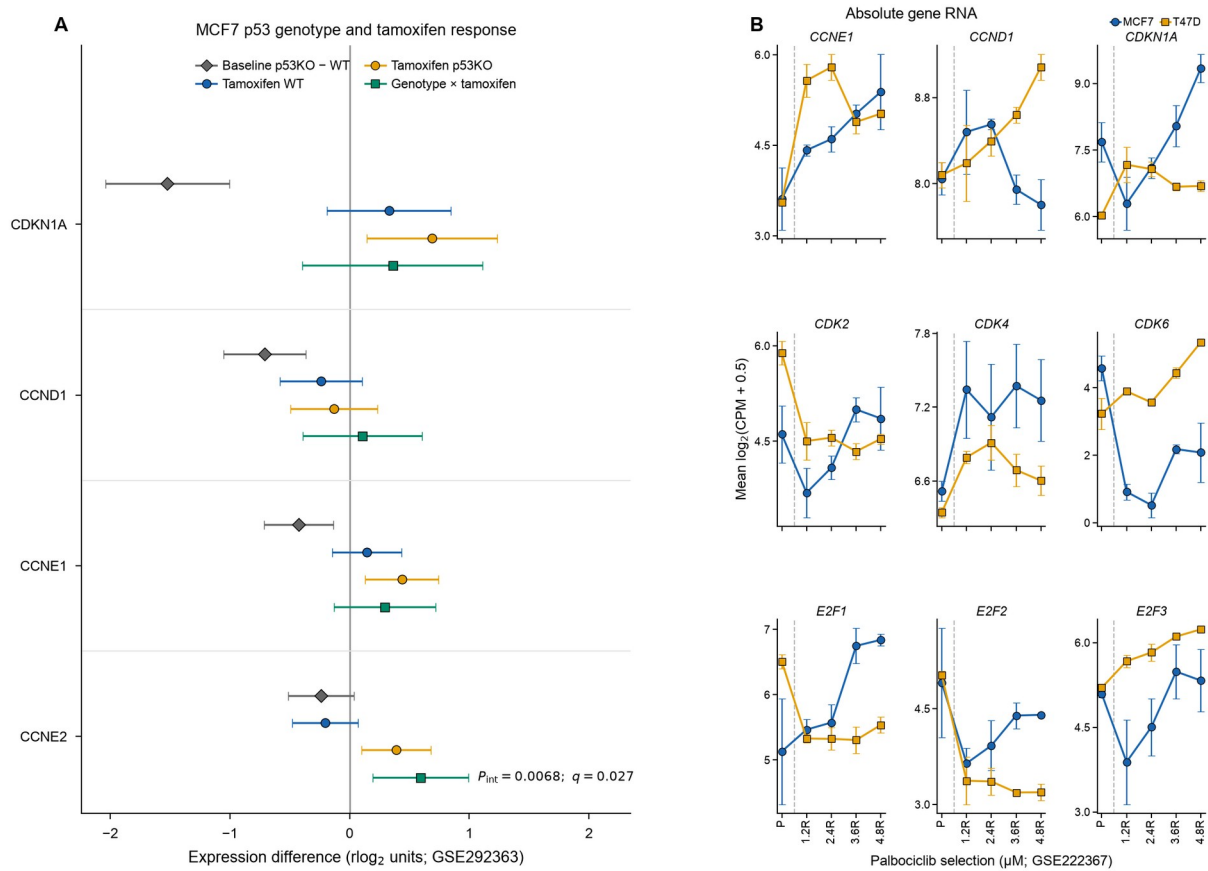

**Fig. S2. p53 genotype, tamoxifen response and palbociclib-resistant cell-cycle states. (A)** OLS contrasts and 95% confidence intervals for MCF7 p53 genotype, tamoxifen and their interaction (GSE292363); $q$ values use Benjamini-Hochberg correction across four genes. **(B)** Mean log<sub>2</sub>(CPM + 0.5) ± s.e.m. for parental and palbociclib-resistant MCF7 and T47D states (GSE222367; $n = 3$ biological samples per state and concentration).

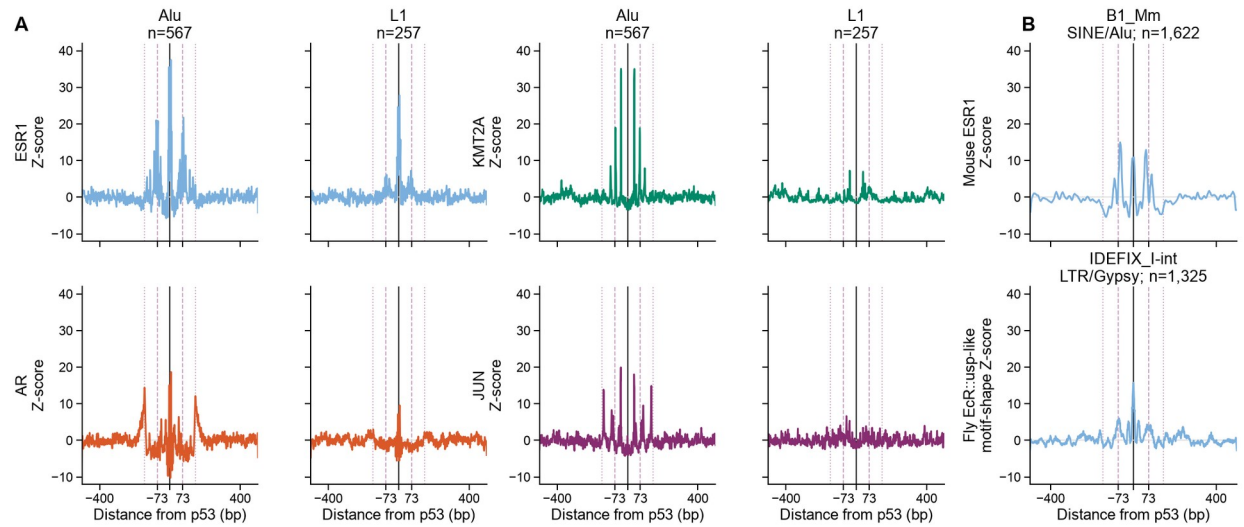

**Fig. S3. Repeat-associated p53-centered motif profiles. (A)** p53-centered ESR1, AR, KMT2A and JUN motif profiles in nutlin-treated MCF10A p53 ChIP-seq peaks overlapping Alu (n = 567) or L1 (n = 257); guides mark 0, +/-73 and +/-146 bp. **(B)** ESR1 motif profiles in mouse B1\_Mm SINEs (n = 1,622) and Drosophila IDEFIX\_I-int LTR/Gypsy elements (n = 1,325).
